# Distinct medial septal GABAergic pathways differentially regulate spatial memory and network activity

**DOI:** 10.64898/2026.11.01.763440

**Authors:** Duncan Archibald Allan MacLaren, Khalid Magomedovich Saidov, Magdalene Isabell Schlesiger, Hannah Monyer

## Abstract

Septal GABAergic projection neurons form distinct pathways to subregions of the hippocampal formation, including the hippocampus proper and the medial and lateral entorhinal cortices. Parvalbumin (PV^+^) and calbindin expressing (CB^+^) projections target different types of interneurons and thereby differentially regulate neuronal activity in the downstream area. Consistent with the idea of dissociable functional roles, inhibition of septal PV^+^ neurons impairs reference memory whereas inhibition of septal CB^+^ neurons impairs both reference and working memory. Here, we show that these functions are differentially supported by specific projection pathways: PV^+^ projections to the dorsal hippocampus and medial entorhinal cortex (MEC) support reference memory, whereas CB^+^ projections to the MEC support both reference and working memory. At the network level, septal PV^+^, but not CB^+^, neurons regulate theta oscillations. Together, these findings further define the roles of septal PV^+^ and CB^+^ pathways in orchestrating network activity and supporting specific forms of memory.

## Introduction

Despite extensive evidence that MS GABAergic signalling regulates hippocampal and entorhinal network dynamics (Unal et al., 2018; Robinson et al., 2023) and behaviour (Quirk et al., 2021; Zheng et al., 2026), how distinct septal GABAergic pathways contribute to spatial memory through different projection targets remains unclear.

Spatial memory enables the learning and retrieval of information about an environment and its features, supporting goal-directed navigation. The hippocampus, particularly the dorsal hippocampus (dHC), plays a central role in spatial memory (Morris et al., 1982; Eichenbaum, 2017), with activity encoding location, trajectory and environmental features (O’Keefe and Dostrovsky, 1971; O’Keefe and Krupic, 2021). The hippocampus is embedded within a broad network of cortical and subcortical structures that provide sensory, contextual and spatial information and regulate network dynamics (Burwell, 2000; van Strien et al., 2009; Hernandez-Frausto and Vivar, 2024). Of particular importance, the medial and lateral entorhinal cortices (MEC and LEC) form major cortical interfaces with the hippocampal formation (Canto et al., 2008; Witter et al., 2017).

The MEC and LEC provide partially distinct but complementary information to the hippocampus. The MEC encodes spatial and navigational information, with neuronal populations exhibiting grid, border and head-direction firing patterns, as well as representations of environmental features including spatial relationships to objects (Fyhn et al., 2004; Hoydal et al., 2019; Caputi et al., 2022; Tukker et al., 2022). In contrast, LEC activity is more strongly associated with object identity and contextual information, although spatially responsive neurons are also present (Deshmukh and Knierim, 2011; Huang et al., 2023). Integration of MEC and LEC inputs may therefore link spatial location with the identity and contextual significance of environmental features (Keene et al., 2016; Connor and Knierim, 2017). Mechanisms coordinating activity across the dHC, MEC and LEC are consequently well positioned to regulate memory.

The medial septum (MS) is a key structure involved in coordinating activity across the hippocampal-entorhinal network (Muller and Remy, 2018; Robinson et al., 2023). GABAergic projection neurons within the MS are heterogeneous, including populations expressing PV or CB (Roland and Savage, 2009; Salib et al., 2019). MS GABAergic neurons play a critical role in the generation of hippocampal theta oscillations (Gerashchenko et al., 2001; Robinson et al., 2024), with MEC gird cell periodicity and accurate performance of memory tasks linked not only to the presence of these oscillations but also their precise timing (Koenig et al., 2011; Quirk et al., 2021; Robinson et al., 2024). Furthermore, theta oscillations are thought to provide a temporal framework that facilitates synchronization between hippocampal, entorhinal, and prefrontal regions, thereby supporting functional interactions during memory processing and decision-making (Colgin, 2011; Tavares and Tort, 2022). Although MS PV^+^ and CB^+^ populations target interneurons, they differ in their postsynaptic targets (Freund and Antal, 1988; Gonzalez-Sulser et al., 2014; Unal et al., 2015). In the MEC, MS PV^+^ inputs preferentially target fast-spiking/PV^+^ interneurons, whereas CB^+^ inputs preferentially engage low threshold spiking (presumed SOM^+^) interneurons (Fuchs et al., 2016). Consistent with these different input targets, activation of MS PV^+^ terminals in the MEC alters theta rhythmic firing, whereas activation of CB^+^ terminals increases burst firing and alters temporal coding in a cell-type specific manner (Schlesiger et al., 2021). We have also shown that inactivation of MS PV^+^ and CB^+^ neurons produces dissociable effects on memory: PV^+^ inactivation impairs reference and contextual memory, whereas CB^+^ inactivation impairs reference and working memory but not contextual memory (Schlesiger et al., 2021). Together, these findings suggest that MS PV^+^ and CB^+^ populations make dissociable contributions to memory.

However, manipulating MS neurons cannot distinguish between behavioural effects mediated through different projection targets. Here, we investigated the contribution of MS PV^+^ and CB^+^ projections to spatial memory by selectively chemogenetically inhibiting their terminals within the dHC, MEC and LEC during a memory-based navigational task. We then investigated whether inhibition of MS PV^+^ or CB^+^ neurons altered LFP theta activity in the MEC. Together, these experiments allowed us to determine whether the contributions of MS GABAergic signalling to memory vary according to source cell type and projection target, and whether these effects are accompanied by changes in MEC LFP theta activity.

## Results

To determine the contributions of MS PV^+^ and CB^+^ projections to the dHC, MEC, and LEC during spatial memory performance, we used pathway-specific chemogenetic inhibition. Inhibitory hM4Di receptors were expressed selectively in MS PV^+^ or CB^+^ neurons, and guide cannulae were implanted bilaterally into the dHC, MEC, or LEC (in separate cohorts of mice) to permit local infusion of CNO. (Figure 1A-C, see S1 for histological analysis). After recovery from surgery, mice were trained in an eight-arm radial maze task which allows reference and working memory to be assessed separately (Figure 1D) (Schmitt et al., 2003) and has previously revealed differential effects of inhibiting MS PV^+^ and CB^+^ neurons (Schlesiger et al., 2021). After task acquisition, we compared the effects of systemic CNO administration with those of local CNO infusion within the target region.

**Figure 1.**
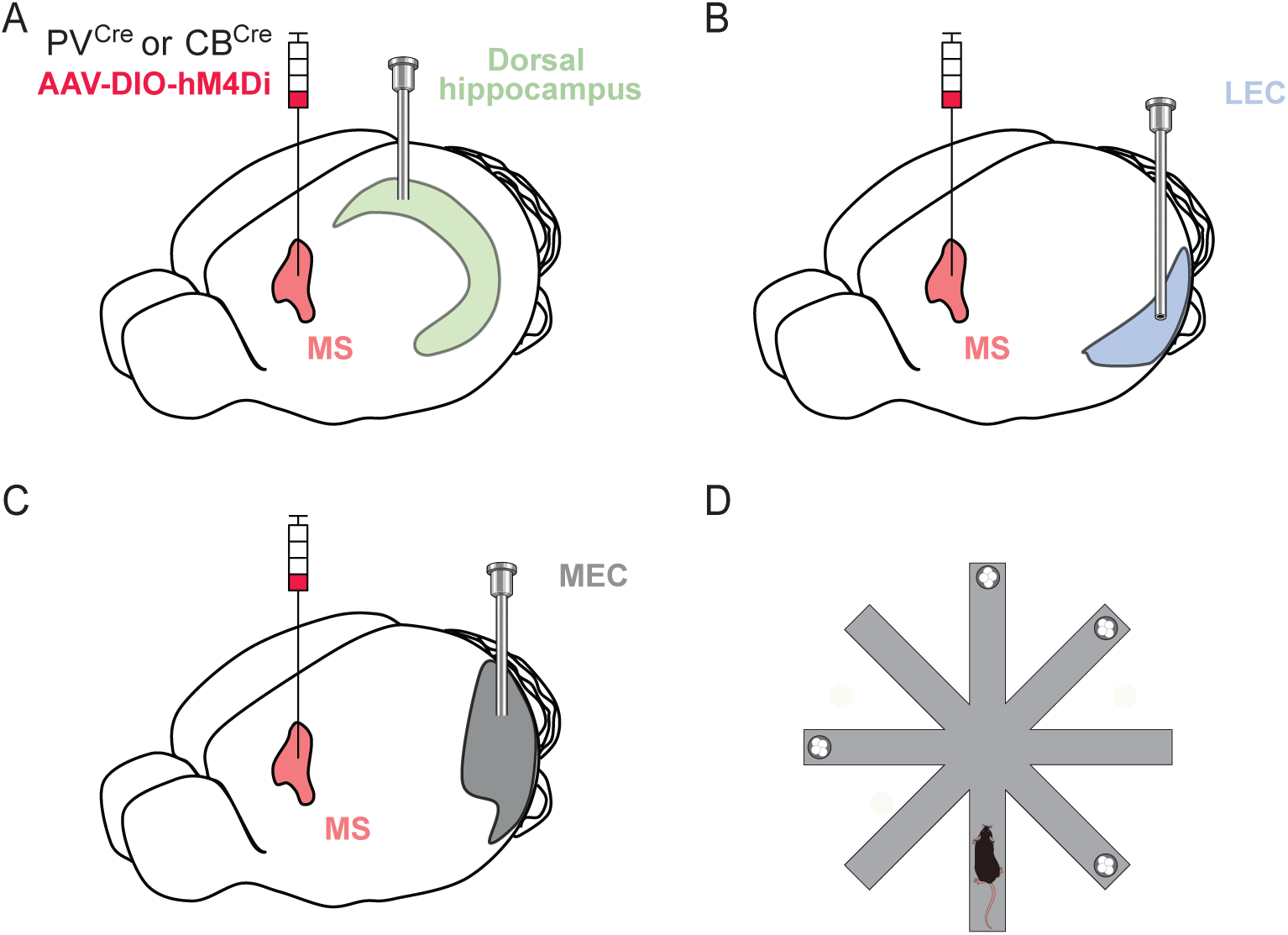
Experimental design for pathway-specific inactivation and spatial memory assessment (A-C) PV^Cre^ and CB^Cre^ mice were injected with AAV-DIO-HM4Di-mCherry into the MS, and guide cannulae were implanted bilaterally into either the dHC, MEC, or LEC to enable pathway-specific inactivation by infusion of CNO into the respective target region. (D) Mice were trained on an 8-arm radial maze task, in which 4 of the 8 arms were baited with reward pellets. During each trial, mice could make two main types of errors: reference memory errors, defined as visits to arms that never contained a reward, and working memory errors, defined as revisits to arms from which the reward had already been collected within that trial.

In PV^Cre^ mice, systemic CNO administration increased reference memory errors per trial but did not affect working memory errors (Figure 2A–F, left). We therefore examined whether this effect was mediated by PV^+^ projections to the dHC, MEC, or LEC. Projection-specific inhibition increased reference memory errors following CNO infusion into the dHC or MEC, whereas inhibition of the LEC projection had no effect (Figure 2A–C, right). Consistent with the lack of an effect of systemic CNO on working memory, inhibition of PV^+^ projections to the dHC, MEC, or LEC did not alter working memory errors (Figure 2D–F, right). There were no increases in reference repeat errors during inhibition of any pathway (Figure S2).

**Figure 2.**
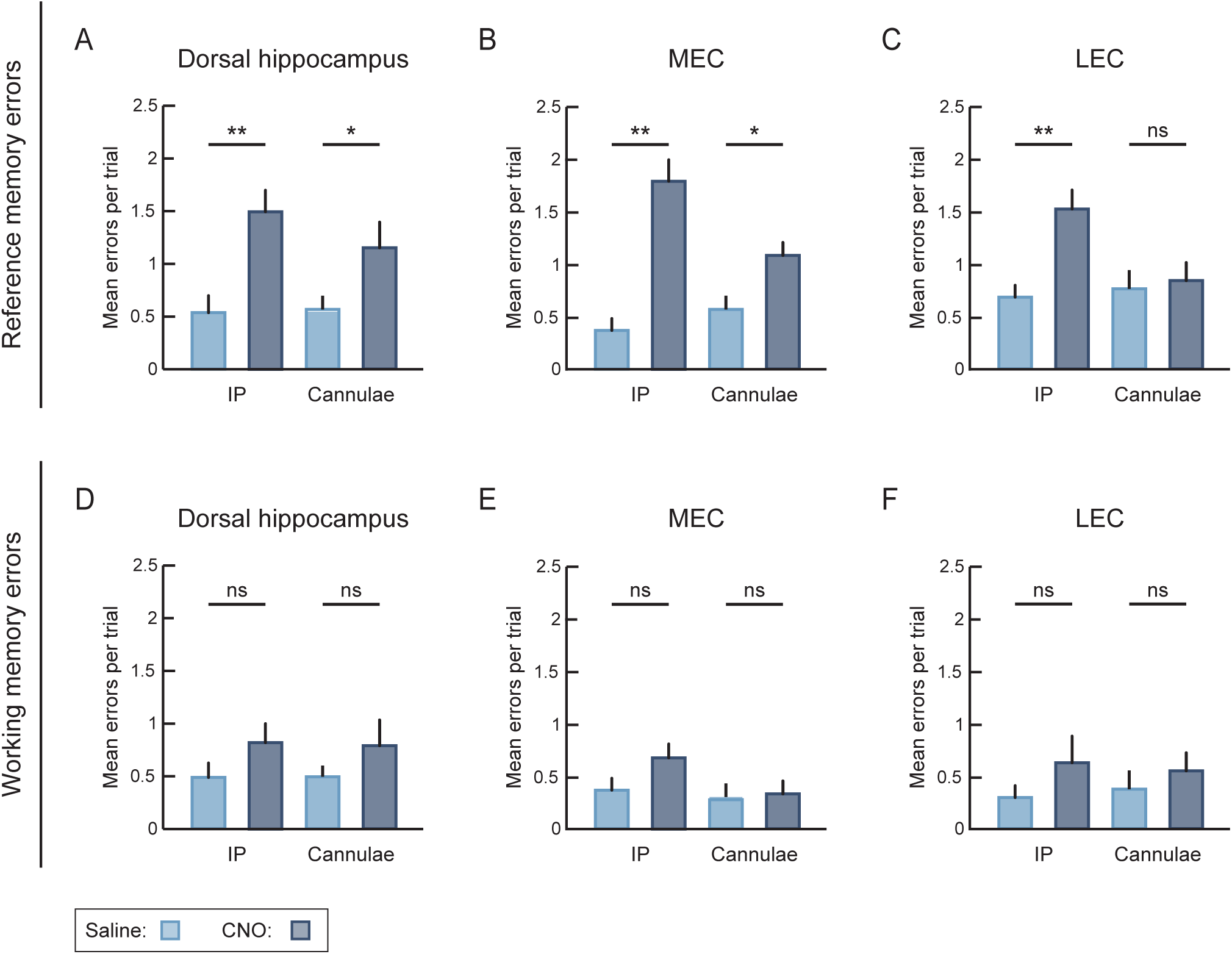
Inhibition of septal PV^+^ projections to the dHC and MEC impairs reference but not working memory. (A–C) Reference memory errors following systemic inhibition of MS PV^+^ neurons or projection-specific inhibition of MS PV^+^ terminals. (A) dHC: systemic CNO administration increased reference memory errors per trial (saline vs CNO, p < 0.01), as did local CNO infusion into the dHC (p < 0.05). (B) MEC: systemic CNO administration increased reference memory errors per trial (p < 0.01), as did local CNO infusion into the MEC (p < 0.05). (C) LEC: systemic CNO administration increased reference memory errors per trial (p < 0.01), whereas local CNO infusion into the LEC had no effect (p > 0.05). (D–F) Working memory errors following systemic inhibition of MS PV^+^ neurons or projection-specific inhibition of MS PV^+^ terminals. (D) dHC: neither systemic CNO administration nor local CNO infusion affected working memory errors (p > 0.05). (E) MEC: neither systemic CNO administration nor local CNO infusion affected working memory errors (p > 0.05). (F) LEC: neither systemic CNO administration nor local CNO infusion affected working memory errors (p > 0.05). All data were analyzed using a mixed- effects model followed by Tukey’s multiple-comparisons test. Data are presented as mean ± SEM. dHC, n = 14 mice; MEC, n = 14 mice; LEC, n = 17 mice. * p < 0.05, ** p < 0.01; ns, not significant.

In CB^Cre^ mice, systemic CNO administration significantly increased both reference and working memory errors per trial (Figure 3A–F, left). Inhibition of CB^+^ projections to the MEC, but not the dHC or LEC, increased reference memory errors (Figure 3A–C, right). Likewise, inhibition of CB^+^ projections to the MEC, but not the dHC or LEC, increased working memory errors (Figure 3D–F, right). There were no increases in reference repeat errors during inhibition of any pathway (Figure S2).

**Figure 3.**
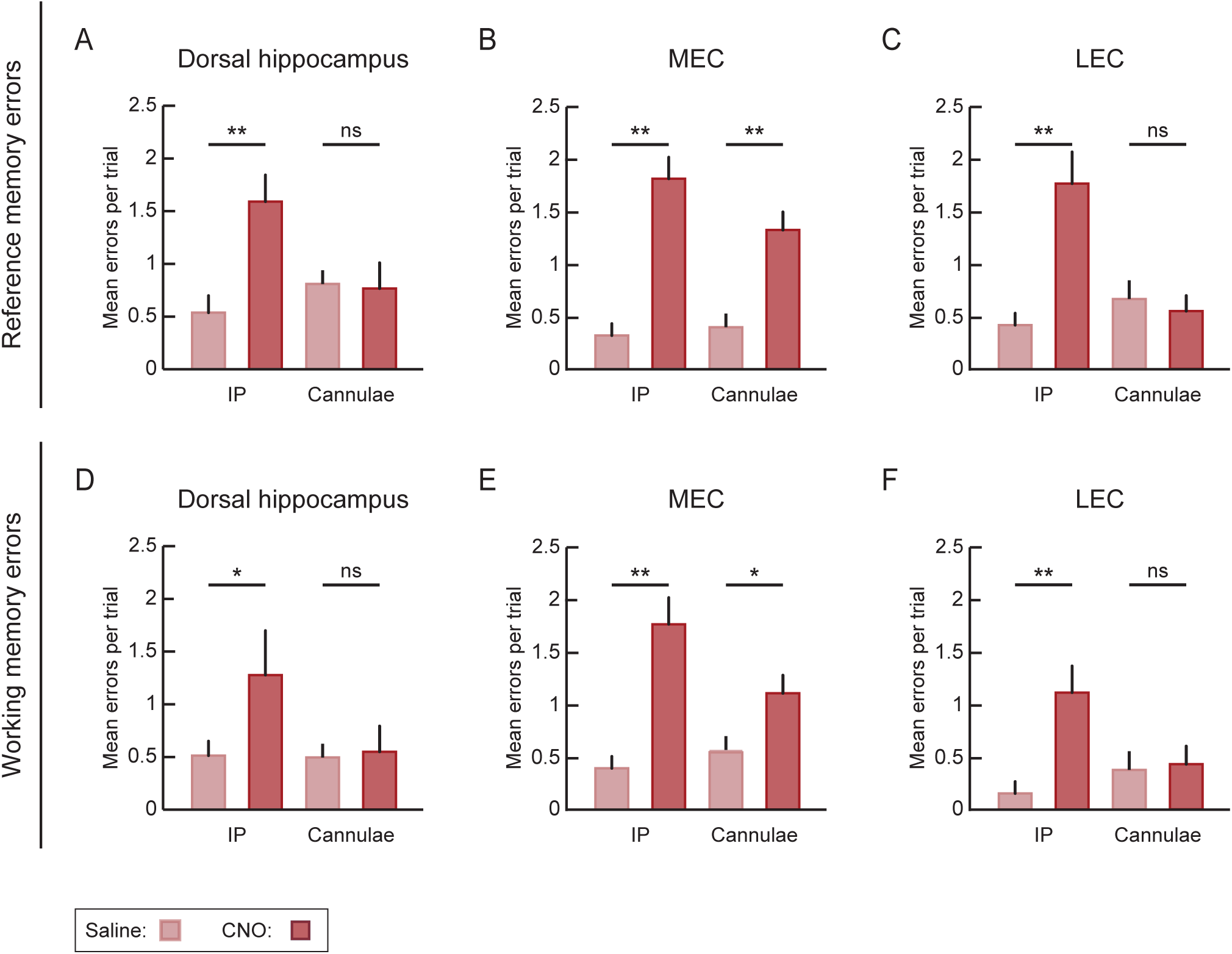
Inhibition of septal CB^+^ projections to the MEC impairs reference and working memory. (A–C) Reference memory errors following systemic inhibition of MS CB^+^ neurons or projection-specific inhibition of septal CB^+^ terminals. (A) dHC: systemic CNO administration increased reference memory errors per trial (saline vs CNO, p < 0.01), whereas local CNO infusion into the dHC had no effect (p > 0.05). (B) MEC: systemic CNO administration increased reference memory errors per trial (p < 0.01), as did local CNO infusion into the MEC (p < 0.01). (C) LEC: systemic CNO administration increased reference memory errors per trial (p < 0.01), whereas local CNO infusion into the LEC had no effect (p > 0.05). (D–F) Working memory errors following systemic or projection-specific inhibition of MS CB^+^ neurons. (D) dHC: systemic CNO administration increased working memory errors per trial (p < 0.05), whereas local CNO infusion into the dHC had no effect (p > 0.05). (E) MEC: systemic CNO administration increased working memory errors per trial (p < 0.01), as did local CNO infusion into the MEC (p < 0.05). (F) LEC: systemic CNO administration increased working memory errors per trial (p < 0.05), whereas local CNO infusion into the LEC had no effect (p > 0.05). All data were analyzed using a mixed-effects model followed by Tukey’s multiple-comparisons test. dHC n = 11 mice, MEC n = 18 mice, LEC = 10 mice. Data are presented as mean ± SEM. * p < 0.05, ** p < 0.01; ns, not significant.

In order to determine whether the memory deficits could be correlated with disruptions to LFP theta oscillations, the presence and dynamics of which have been implicated in memory (Quirk et al., 2021; Etter et al., 2023), we recorded LFP activity in the MEC during inactivation of MS PV^+^ or CB^+^ neurons (Figure 4A). Inactivation of MS PV^+^ neurons reduced MEC theta power without altering theta frequency (Figure 4B, 4D). In contrast, inactivation of MS CB^+^ neurons produced no detectable changes in either the frequency or power of MEC theta oscillations (Figure 4C, 4E).

**Figure 4.**
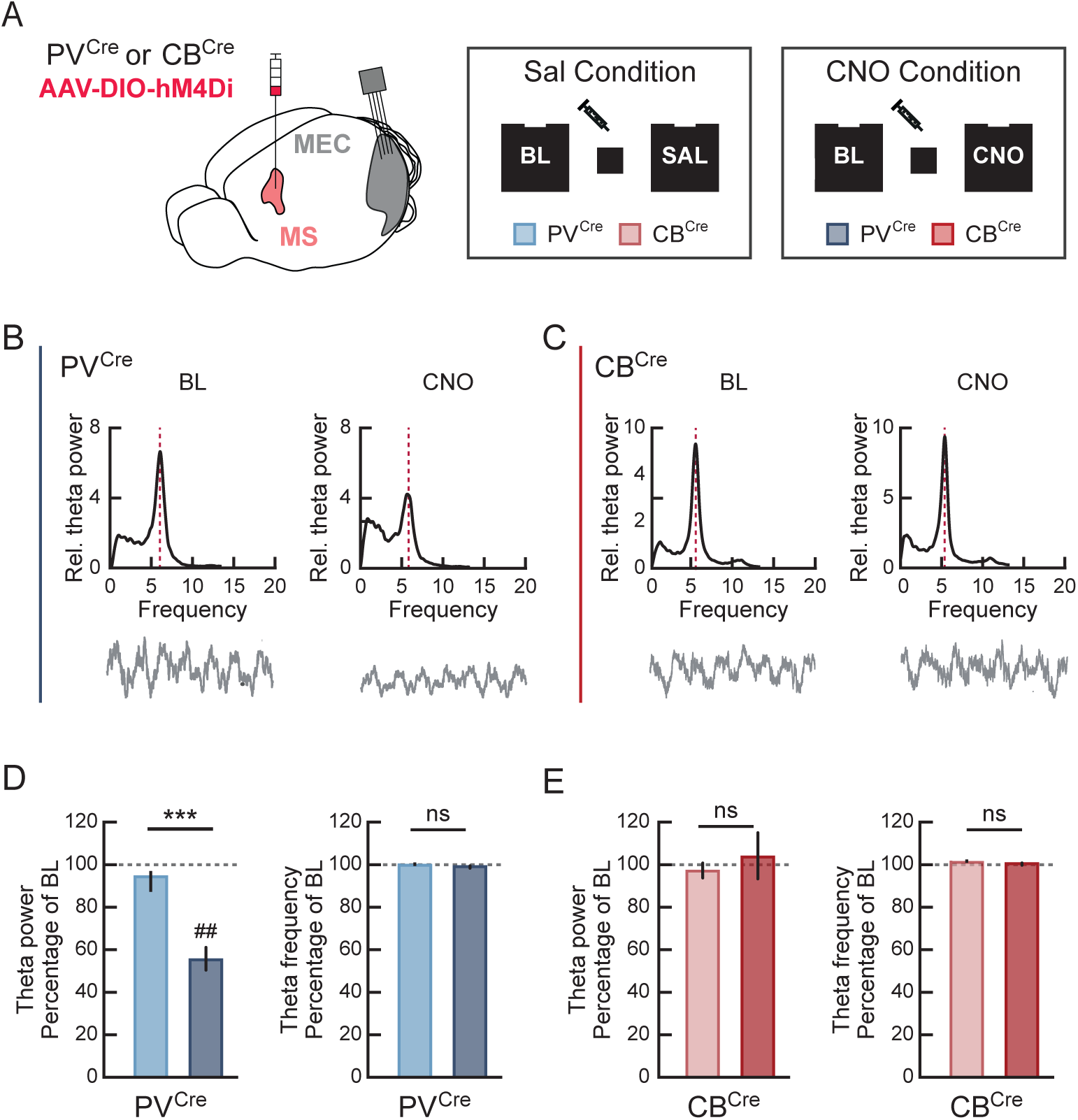
Inactivation of MS PV^+^ but not CB^+^ neurons reduces LFP theta oscillations in the MEC. (A) AAV-DIO-hM4D(Gi)-mCherry was injected into the MS of PV^Cre^ or CB^Cre^ mice, and LFP recordings were performed in the MEC before and after intraperitoneal injection of saline (SAL) or CNO. (B) Power spectral density (PSD) plots of LFP activity (0–20 Hz) recorded from a representative tetrode in a PV^Cre^ mouse during the indicated conditions. Stippled lines indicate the peak frequency within the theta range. A representative 1-s raw LFP trace is shown below. (C) Same as in (B), but from a CB^Cre^ mouse. (D) CNO administration decreased LFP theta power in PV^Cre^ mice (SAL vs. CNO p ≤ 0.001; CNO vs. 100% of baseline p ≤ 0.005), but did not alter theta frequency (all p > 0.05). (E) CNO administration in CB^Cre^ mice had no effect on theta power or frequency (all p > 0.05). SAL v CNO comparisons were analyzed using the Mann–Whitney U test and comparisons with 100% of baseline were analyzed using the Wilcoxon signed-rank test. PV^Cre^, n = 5 mice; CB^Cre^, n = 6 mice. Bar graphs show mean ± SEM. ^##^ p ≤ 0.005, *** p ≤ 0.001.

These findings demonstrate region and cell type-dependent contributions of MS GABAergic projections to spatial memory and LFP theta generation. PV^+^ projections to the dHC and MEC contribute to reference memory and the generation of LFP theta, whereas CB^+^ projections selectively to the MEC support working and reference memory without contributing to LFP theta generation.

## Discussion

The present findings demonstrate that MS GABAergic regulation of spatial memory and LFP theta oscillations is organized according to both neuronal identity and projection target. Pathway-specific inhibition revealed that PV+ inputs to the dHC and MEC and CB+ inputs to the MEC contribute to reference memory, whereas only inhibition of the CB+ to MEC pathway impaired working memory. Neither cell type influenced memory through projections to the LEC. These results identify specific downstream pathways through which MS PV+ and CB+ neurons make dissociable contributions to distinct components of spatial memory.

The effects of MS PV^+^ pathway inhibition suggest that MS PV^+^ signaling contributes to reference memory on the 8-arm radial maze through both hippocampal and medial entorhinal targets. This is consistent with the established roles of these regions in spatial memory (Hales et al., 2014; Eichenbaum, 2017) and of MS PV^+^ input in regulating network activity, particularly the theta rhythm (Quirk et al., 2021; Robinson et al., 2023). Theta oscillations have been linked to MEC grid-cell periodicity (Koenig et al., 2011; Robinson et al., 2024), and accurate performance of delayed spatial alternation tasks has been attributed to their presence (Etter et al., 2023). Thus, the impairment following MS PV^+^ pathway inhibition to the dHC and MEC may reflect, at least in part, disrupted theta-dependent coordination within hippocampal and entorhinal networks.

In contrast, MS CB^+^ inactivation also produced target-specific effects on radial-maze performance, with selective inhibition of the CB^+^ projection to the MEC impairing both reference and working memory. Importantly, unlike MS PV^+^ inactivation, which reduced MEC theta power, MS CB^+^ inactivation produced no detectable changes in LFP theta activity. Thus, despite the similar effects of inactivation of the MS PV^+^ and CB^+^ projections to the MEC on reference memory, and the additional impairment of working memory following CB^+^ pathway inactivation, the memory effects of CB^+^ inactivation are dissociable from changes in MEC theta activity.

A potential explanation for the distinct effects of PV^+^ and CB^+^ pathway inactivation lies in their differential recruitment of local inhibitory circuits. Although both MS projections exclusively target interneurons, PV^+^ projections preferentially target PV^+^ interneurons, whereas CB^+^ projections preferentially target LTS (presumed SOM^+^) interneurons (Fuchs et al., 2016). These interneuron populations have distinct patterns of connectivity within the local network (Tukker et al., 2022) and differentially regulate spatial coding: PV^+^ interneuron inactivation disrupts grid firing and speed coding, whereas SOM^+^ interneuron inactivation reduces spatial selectivity (Miao et al., 2017). Further evidence that MS PV^+^ and CB^+^ inputs differentially influence MEC network dynamics comes from our previous work, showing that MS PV^+^ input regulates theta activity, whereas activation of CB^+^ terminals increases burst firing and enhances phase precession in a cell-type-specific manner (Schlesiger et al., 2021). Together, these findings raise the possibility that the selective impairment of working memory following MS CB^+^ terminal inactivation in the MEC reflects disrupted temporal coding in MEC spatial cells, potentially compromising the maintenance or updating of spatial information during ongoing task performance.

The absence of behavioral effects following inhibition of MS projections to the LEC is also informative. The LEC is an important component of the hippocampal–entorhinal network and contributes to the representation of object identity, contexts, spatial, and object-context information (Deshmukh and Knierim, 2011; Huang et al., 2023). Given the sparse PV^+^ projection to the LEC and comparatively low theta activity within this region (Deshmukh et al., 2010), the absence of an effect following PV^+^ pathway inhibition is perhaps not surprising. However, MS CB^+^ projections to the LEC are considerably denser than PV^+^ projections (unpublished observations), and given the effect of CB^+^ terminal inhibition in the MEC on working memory in this task, the lack of an effect following CB^+^ terminal inhibition in the LEC points to an additional functional distinction between these two entorhinal regions.

Individual MS GABAergic neurons can collateralize to multiple downstream regions (Fuchs et al., 2016; Viney et al., 2018), suggesting that these pathways may influence network function as part of broad, distributed circuits. However, our findings support a model in which MS GABAergic neurons do not exert a uniform influence over hippocampal-entorhinal function, but instead coordinate network activity through anatomically and functionally differentiated pathways. MS PV^+^ projections to the dHC and MEC contribute to reference memory and regulate LFP theta, whereas MS CB^+^ projections to the MEC contribute to both reference and working memory without contributing to LFP theta activity. This dissociation indicates that the contribution of MS GABAergic pathways to spatial memory is determined by both the identity of the projecting neuron and its downstream targets.

## Materials and methods

Experiments were performed on naive adult male heterozygous PV^Cre^ (Hippenmeyer et al., 2005) and CB^Cre^ (Fuchs et al., 2016) mice that were maintained on a C57BL/6N background. Mice were 14-20 weeks old at the time of surgery. Mice were housed on a 12 h light/dark schedule, and the experiments were performed during the light phase. All procedures were approved by institutional animal care committees and by the governmental supervisory panel on animal experiments of Baden Wurttemberg in Karlsruhe and conduced in compliance with the European Council Directive 86/609/EEC.

## Surgical procedures

Mice were anesthetized with isoflurane and mounted in a stereotaxic frame where isoflurane anesthesia (1.0–2.5%) was maintained via a facemask. The skull was exposed and a craniotomy was made above the MS and 400 nL of hSyn-DIO-hM4D(Gi)-mCherry (44362- AAV8, Addgene) injected from a glass pipette, the first 200 nL at AP: +1.00 mm; ML: ± 0.0 mm; DV: -4.1, after 300 s additional 200 nL were injected at AP: +1.00; ML: ± 0.0; DV: -3.7. The pipette was removed after 300 s. The skull was cleaned and 2-3 miniature screws inserted. For cannulae implantation 26-gauge guide cannula (C315GA/SPC, Plastics One) were implanted bilaterally into one of the three target regions at the following co-ordinates. dHC: AP -2.2; ML ± 2.0; DV -1.5mm and angled 10° towards the midline (the angle was to avoid the dummy cannula colliding). MEC: 0.5mm in front of superior sagittal sinus; ML ± 3.1; DV: -2.5 with a 10° angle posterior. LEC: AP: -3.8, ML: ± 4.3; DV -1.5 angled 4° lateral. Cannula were secured to the screws and skull with dental cement and the wound closed with sutures. A dummy cannula (C315DC/SPC, PlasticsOne) was inserted into each guide cannula. For microdrive implantation, prior to implantation, tetrodes were constructed by twisting tungsten wire (California Fine Wire Company) and subsequently mounted into custom designed 4 tetrode microdrives (Axona) that allowed the individual movement of the tetrodes. A craniotomy was made above the MEC and tetrodes were implanted ∼0.3 mm in front of the medial sinus, +3.1mm from the midline, 0.8mm below dura at an angle of 7° in the posterior direction. The microdrive was attached to the screws and skull with dental cement and the wound closed with sutures. For all surgeries, carprofen was administered on the day of surgery and 2 days post-surgery. Mice were allowed > 7 days to recover before experiments began.

## Radial maze training and testing

The testing apparatus consisted of a standard 8 arm radial maze with 8 arms (28 cm long and 5 cm wide) extending from a central platform (18.5 cm in diameter). Each arm had a small bowl at the end to hold reward pellets. The maze was at an elevation of 40.5 cm from the floor and situated in a dimly lit testing environment containing prominent distal cues. Mice were food restricted to 85%–90% of free food weight prior to the start of training. For each mouse, 4 arms were assigned as baited arms and the remaining 4 arms assigned as non-baited. The baited arms varied between mice, but always had the same pattern (e.g., baited: 1,2,5,7). Mice had 4 trials per day with 60 min between trials. At the start of the trial 1-2 reward pellets (AIN-76A Rodent Tablet 5 mg, TestDiet) were placed in the bowl at the end of each baited arm and the mouse was placed on the central platform. The trial ended after all pellets had been consumed or 5 min had elapsed, after which the maze was cleaned with a mild detergent. Arm entries were classified into 4 types: correct (entry into a baited arm which still contained the reward); working memory error (re-entering an arm from which the reward had already been collected within that trial); reference memory error (entering a never baited arm for the first time in that trial); reference memory repeat error (re-entering a reference memory error arm within the same trial) (Schmitt et al., 2003; Schlesiger et al., 2021). Mice were considered to have learnt the task when all rewards were collected and mean reference memory errors were < 1 per trial, which took upto 30 days. Once these criteria were reached, testing began. Mice were administered saline by intraperitoneal (i.p.) injection (10 ml/kg) and the following day i.p. CNO (3 mg/kg in saline at a volume of 10 ml/kg, Tocris) 20 mins prior to the first trial. For cannulae infusions mice were gently restrained and the dummy cannulae removed, guide cannulae (C315IA/SPC, PlasticsOne) were then inserted which extended 0.8mm beyond the injector tip and 1uL of saline or CNO (either 10µM or one of 0.1 µM, 1.0 µM, and 100 µM in saline) was infused. Infusions were conducted over 60s and cannulae were left in situ for a further 120s before being removed and replaced with dummy cannulae. See Figure S1 for examples of viral expression and cannulae tracks. See Figure S2 for a CNO dose-response on an initial cohorts of mice. Analysis and figures in the main text show results from the 10 µM CNO concentration.

## Electrophysiological recordings

At least one week after surgery, mice were food restricted to ∼90% of their free food weight and trained to forage for sugar pellets (Ain-76A Rodent Tablet 5 mg, TestDiet) randomly distributed by a dispenser (Med Associates) in a black open field arena (70 cm × 70 cm x 30 cm) with a polarizing white cue card (21 cm × 30 cm) on one wall. LFP recordings were performed in two 20-min foraging sessions in the open field with and intertrial interval of 30 min during which time the mouse was placed in a smaller rest box. On two consecutive days, 10 ml/kg of either saline or CNO (3 mg/kg) was injected I.P. immediately before the inter- trial rest box. Electrophysiological signals were amplified and digitized by a 16-channel amplifier board (RHD2132, Intan Technologies) connected by a lightweight cable to a RHD2000 (Intan Technologies) data acquisition board with a sampling frequency of 20 kHz. The position of the mouse in the open field arena was triangulated from the position of three LEDs (red, green, blue) that were mounted on the amplifier headstage. The position of each LED was recorded with a camera (DFK 23GM021, The Imaging Source) using the ’positrack’ software (https://github.com/kevin-allen/positrack). Sampling timestamps of each tracking position were recorded along with the electrophysiological signal using the ‘ktan’ software (https://github.com/kevin-allen/ktan). The average LFP theta amplitude and frequency per mouse was then calculated for baseline and saline/CNO treatment sessions. For each tetrode, a time-frequency spectrogram from 0 to 30 Hz was calculated via the Chronux55 (http://www.chronux.org/) function mtspecgramc() using a window size and time step of 20 s and 10 s, respectively. The LFP theta frequency was taken as the frequency at maximum power within the 6–12-Hz frequency band of the time-averaged spectrogram. The time-averaged spectrogram S(f) was then 1/f-corrected via the normalization 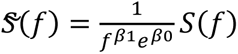 where β0 and β1 are coefficients obtained from the linear regression of log S(f) on log (f). The relative power was then taken as the maximum power of the normalized spectrum in the 6–12-Hz frequency band. Tetrodes were advanced by 50-100 µm after each recording session with good coverage. This procedure yielded 4-8 session pairs per mouse.

## Histology

After completing experiments, mice were deeply anesthetized with ketamine and xylazine, and perfused intracardially with phosphate buffered saline (PBS) followed by 4% paraformaldehyde. Heads were left in 4% paraformaldehyde overnight. Brains were subsequently block cut such that the region including the MS, dHC and LEC (where appropriate) could be further processed in the coronal plane and the region including the MEC further processed in the sagittal plane. Fifty µm sections were cut using a vibratome. For analysis of virus expression at the injection site, the region containing the MS was stained with DAPI and mounted with Mowiol 40–88 on glass slides. The regions containing the cannulae and tetrode tracks were mounted onto glass slides and stained with cresyl violet. Mice without clear virial expression or with cannulae outside the intended target region were removed from all analyses.

## Acknowledgements

We thank Drs A. Caputi and E. Fuchs for interesting discussions throughout the project, Livia Burger for assistance in data collection, and Tahir Mehmood for excellent technical assistance. We thank and acknowledge the staff and services offered by the Light Microscopy Facility at the DKFZ. The project was funded by DFG Research Grants CRC/TRR 1436 and CRC/TRR 384 H.M., Advanced ERC research grant 101142587 to H.M., and DFG Emmy Noether Programme grant 469030888 to M.I.S.

## Author contributions

Conceptualization: H.M., and D.A.A.M.

Methodology and experimentation: K.M.S., M.I.S., and D.A.A.M. Analysis: D.A.A.M., and M.I.S.

Funding acquisition: H.M., and M.I.S. Writing: D.A.A.M. and H.M.

---

**Figure S1.**
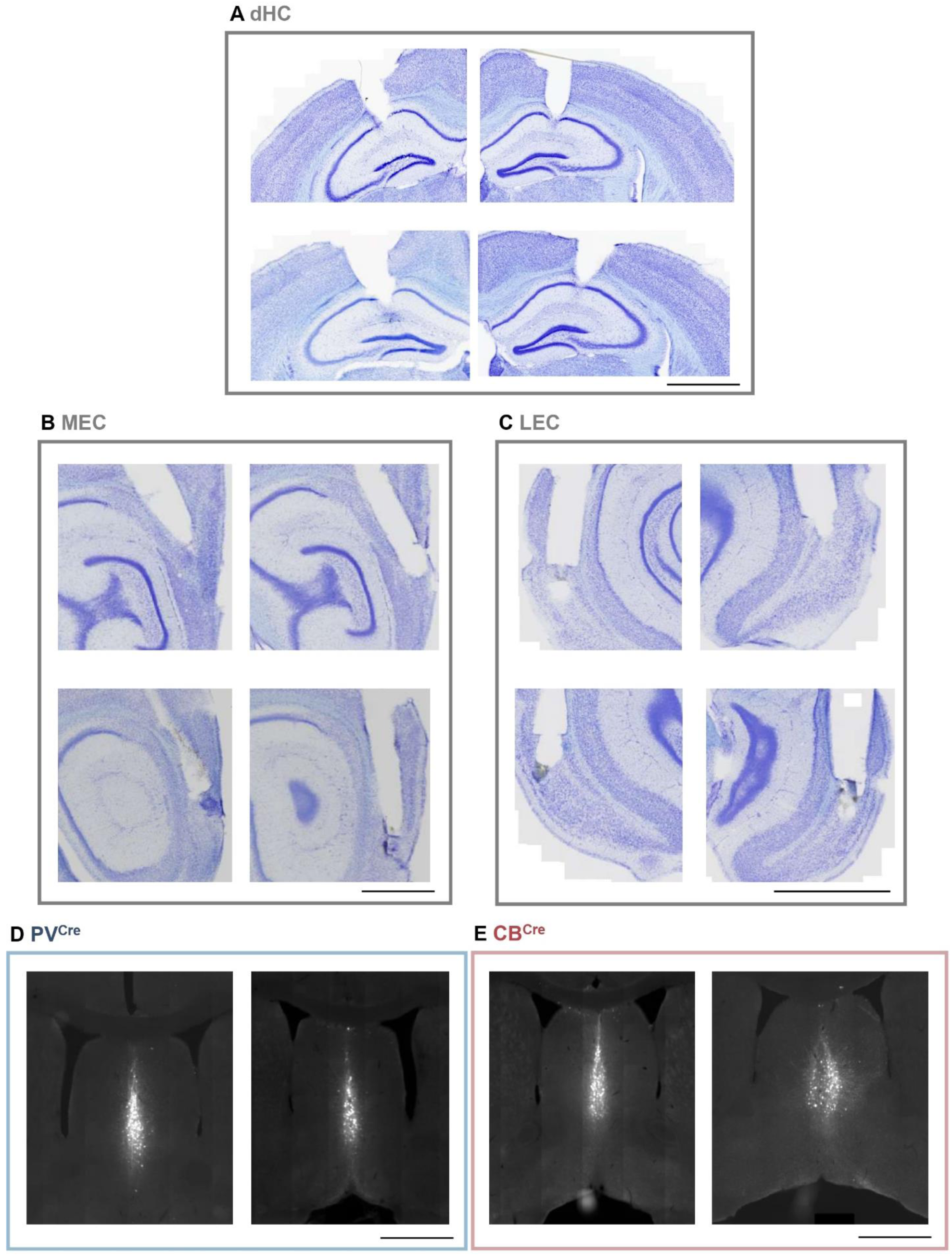
Example implantation sites and viral expression (A-C) Photomicrographs of Nissl-stained coronal (dHC and LEC) and saggital (MEC) sections showing examples of cannula implantation sites. (D-E) mCherry expression (white) in the MS in representative PV^Cre^ (left panels) and CB^Cre^ (right panels) mice. Scale bar indicates 1000 µM.

**Figure S2.**
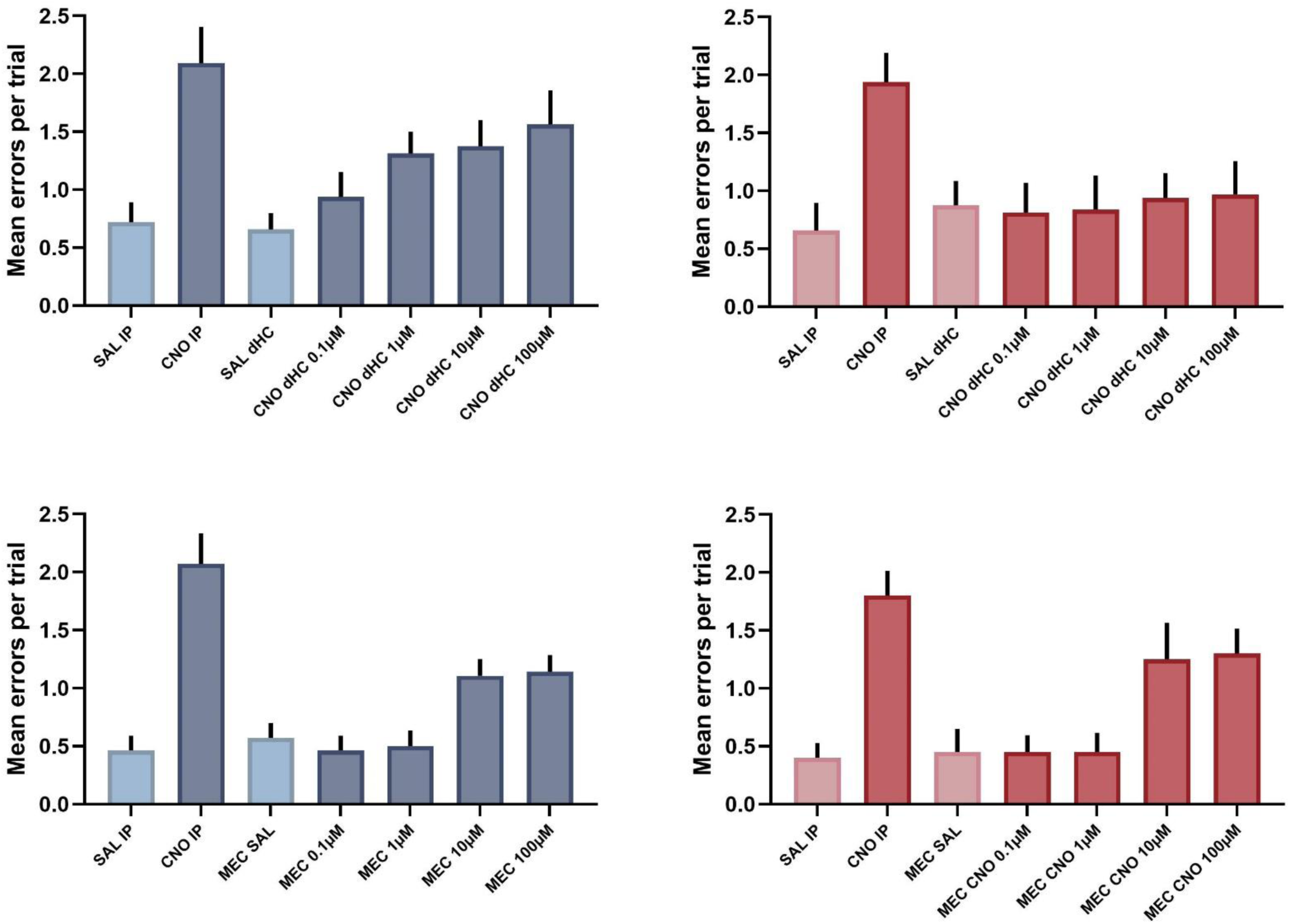
CNO dose-response experiment in initial cohorts of mice used to establish the 10µM concentration used in subsequent experiments and analysis Reference memory errors following systemic CNO or infusions of differing concentrations of CNO into the dHC (top rows) or MEC (bottom rows) in PV^Cre^ mice (left panels) or CB^Cre^ mice (right panels). The combined analysis of these and subsequent cohorts of mice presented in the main text shows the 10µM concentration. PV dHC n = 8 mice, CB dHC n = 8 mice, PV MEC n = 7 mice CB MEC n = 6 mice. Data are presented as mean ± SEM.

**Figure S3.**
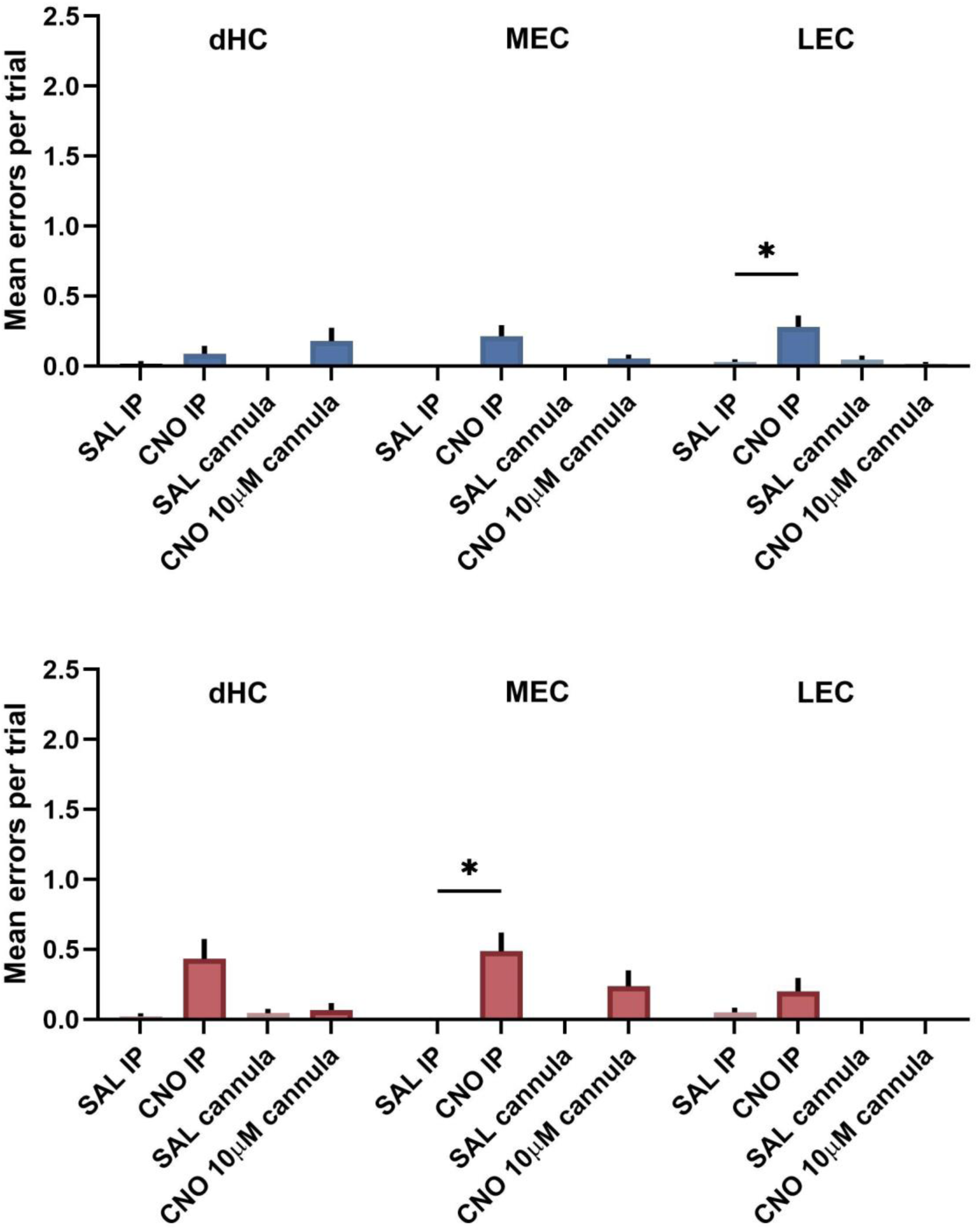
Inhibition of MS PV^+^ or CB^+^ neurons, or their projections, has no systematic effect on the number of reference repeat memory errors Reference repeat errors (also referred to as ‘working memory incorrect errors’ Schmitt et al., 2003) during manipulation of PV^+^ neurons or their projections (top panel) or CB^+^ neurons and their projections (bottom panel) to the indicated regions. Manipulation of either cell type had no systematic effect on this type of error (in line with what we have previously shown for systemic CNO administration, Schlesiger et al. 2021). All data were analyzed using a mixed-effects model followed by Tukey’s multiple-comparisons test. * p < 0.05. Data are presented as mean ± SEM. PV+ dHC, n = 14 mice, MEC, n = 14 mice, LEC, n = 17 mice. CB MEC n = 18 mice, LEC = 10 mice. Data are presented as mean ± SEM.

